# Binding and sequestration of poison frog alkaloids by a plasma globulin

**DOI:** 10.1101/2022.11.22.517437

**Authors:** Aurora Alvarez-Buylla, Maria Dolores Moya-Garzon, Alexandra E. Rangel, Elicio E. Tapia, Julia Tanzo, H. Tom Soh, Luis A. Coloma, Jonathan Z. Long, Lauren A. O’Connell

## Abstract

Alkaloids are important bioactive molecules throughout the natural world, and in many animals they serve as a source of chemical defense against predation. Dendrobatid poison frogs bioaccumulate alkaloids from their diet to make themselves toxic or unpalatable to predators. Despite the proposed roles of plasma proteins as mediators of alkaloid trafficking and bioavailability, the responsible proteins have not been identified. We use chemical approaches to show that a ~50 kDa plasma protein is the principal alkaloid binding molecule in blood from poison frogs. Proteomic and biochemical studies establish this plasma protein to be liver-derived alkaloid-binding globulin (ABG) that is a member of the serine-protease inhibitor (serpin) family. In addition to alkaloid binding activity, ABG sequesters and regulates the bioavailability of “free” plasma alkaloids *in vitro*. Unexpectedly, ABG is not related to saxiphilin, albumin, or other known vitamin carriers, but instead exhibits sequence and structural homology to mammalian hormone carriers and amphibian biliverdin binding proteins. Alkaloid-binding globulin (ABG) represents a new small molecule binding functionality in serpin proteins, a novel mechanism of plasma alkaloid transport in poison frogs, and more broadly points towards serpins acting as tunable scaffolds for small molecule binding and transport across different organisms.

## 1. INTRODUCTION

Alkaloids are nitrogenous small molecules that play important ecological and physiological roles throughout nature, one of which is mediating predator-prey interactions. Species across many taxa, including plants, insects, marine invertebrates, and vertebrates, have co opted alkaloids as chemical defenses, methods for hunting, and pheromone signals. Some of the most potent alkaloid toxins, including batrachotoxin (BTX), saxitoxin (STX), and tetrodotoxin (TTX), act specifically by affecting voltage gated sodium (NaV), potassium, and calcium channels, leading to disruption of nerve and muscle cells [1–3]. In the blue-ringed octopus, *Hapalochlaena lunulata*, TTX is used to paralyze prey [4], while in the pufferfish *Takifugu niphobles* it also acts as a pheromone [5], and in the California newt *Taricha torosa* it is a defense against predation [6]. Other less-potent alkaloids also play important roles in predator-prey interactions. For example, Lepidoptera insects (butterflies and moths) and Coleoptera beetles sequester pyrrolizidine alkaloids from plants for predation, defense, and production of pheromones [7,8]. Although the identities of these alkaloids are well documented, less is known about the physiological mechanisms that allow animals to produce, sequester, and autoresist these small molecules. Identifying and characterizing proteins that interact with alkaloids in these ecological contexts allows us to better understand how animal physiology has coevolved with alkaloids.

Despite the important ecological and physiological roles of alkaloids in animals, the molecular mechanisms involved in alkaloid production, transport, and resistance have been more elusive and typically focused on a single alkaloid or specific structural class of alkaloids. In grasshoppers and moths, passive absorption of pyrrolizidine alkaloids is followed by conversion into a non-toxic form by haemolymph flavin-dependent monooxygenase, allowing the insects to avoid autointoxication [9]. In some beetle species, ATP-binding cassette (ABC) transporters actively pump pyrrolizidine alkaloids into reservoir defensive glands [10]. In vertebrates, the proteins that allow for the sequestration of alkaloids without autotoxicity are unclear with the exception of previous work with TTX and STX. The pufferfish saxitoxin and tetrodotoxin binding protein (PSTBP) was originally identified in the plasma of *Fugu pardalis [11]*, and is thought to play a role in the transport of TTX and STX to the site of bioaccumulation in the liver and ovaries in many pufferfish species [12]. The soluble protein saxiphilin has been proposed as a toxin sponge for STX in various species of amphibians [13], although it remains unclear whether these species come into contact with STX in nature and whether saxiphilin may act as the predominant STX transporter in the plasma of frogs. While these insights have advanced our understanding of toxin physiology, studies in vertebrates have been narrowly focused on a few potent alkaloids with high-specificity binding proteins. However, other animals carry a wide diversity of alkaloids for chemical defense and the physiological mechanisms that allow accumulation of structurally diverse alkaloids are unknown.

Some species of frogs sequester a remarkable diversity of dietary alkaloids onto their skin as a chemical defense. This trait has independently evolved in several frog families, including Dendrobatidae in Central and South American and Mantellidae in Madagascar. Over 500 compounds have been found on the skin of *Dendrobatidae* frogs, with some alkaloids sourced from ants, mites, millipedes and beetles [14–16]. Within dendrobatids, alkaloid-based chemical defenses have evolved independently at least three times [17,18], where non-toxic species do not uptake alkaloids onto their skin even when they are present in their diet [19–22]. Well-studied poison frog alkaloids include pumiliotoxins (PTX), whose documented targets include sodium and potassium gated ion channels [23,24], and decahydroquinolines (DHQ), which affect nicotinic acetylcholine receptors [25]. Epibatidine was first identified in the genus *Epipedobates* and specifically binds certain nicotinic receptors, leading it to be proposed as a therapeutic analgesic alternative to morphine [26]. Although there is limited research into the mechanisms of sequestration and autoresistance of alkaloids in poison frogs [13,27–30], it is likely this process involves alkaloid transport through circulation for these dietary compounds to end up in skin storage glands. Based on the extensive work on plasma small molecule transport in mammals, one might expect that proteins like albumin, which is an abundant and promiscuous small molecule binder in the blood [31–33], or vitamin transporters [34–36], which are able to interact with diet derived molecules, might be involved in alkaloid sequestration in poison frogs. In this study, we tested the hypothesis that poison frogs have an alkaloid-binding protein in the plasma and aimed to uncover its functional role and evolutionary significance. We predicted this protein would bind a range of poison frog alkaloids and would be present in frogs that are chemically defended in nature, but not in undefended species.

## 2. RESULTS

### An alkaloid-like photoprobe identifies a binding protein in poison frog plasma

We used a biochemical strategy to directly test for alkaloid binding in the plasma of different species of poison frogs. To do this, we obtained a UV crosslinking probe with an indolizidine functional group that shares structural similarity to the poison frog alkaloid pumiliotoxin **251D** (PTX) (Figure 1A, functional group highlighted in blue). Upon UV irradiation, the diazirine (green) enables protein crosslinking and the subsequent probe-protein complex can be conjugated to a fluorophore for gel-based visual analysis or biotin for streptavidin enrichment. Application of this photocrosslinking approach outside of mammalian systems has been remarkably limited, and in frogs has been limited to studying neuromuscular receptors [37,38]. We found the PTX-like photoprobe shows binding activity within the plasma of three species of dendrobatid poison frogs, *Oophaga sylvatica, Dendrobates tinctorius*, and *Epipedobates tricolor* (Figure 1B). In these species, this binding activity was largely restricted to a few bands in the 50-60 kDa range, and was a similar size across different species. Plasma from a non-toxic dendrobatid poison frog (*Allobates femoralis*), a mantellid poison frog (*Mantella aurantiaca*), the cane toad (*Rhinella marina*), and humans showed no binding activity with the photoprobe (Figure 1B). We further tested whether the presence of alkaloids would compete off photoprobe binding. In *O. sylvatica*, photoprobe binding resulted in two bands that showed competition by the addition of PTX, decahydroquinoline (DHQ), or epibatidine (epi), but not with nicotine (Figure 1C). In *D. tinctorius*, the photoprobe showed a two-band binding pattern, where the bottom band was competed by PTX and there was slight competition of both bands with DHQ and epibatidine, but no competition with nicotine (Figure 1D). In both *O. sylvatica* and *D. tinctorius*, competition occurred when PTX was 10-fold higher in concentration than the photoprobe (Figure S1A,C). In *E. tricolor* plasma from some individuals two bands were observed while in others only one band was seen, and these were more faint in the presence of DHQ or epibatidine but not PTX or nicotine (Figure 1E). We conclude from these photocrosslinking experiments that plasma binding of alkaloids in three species of chemically defended dendrobatid poison frogs is mediated by a ~50-60 kDa plasma protein.

**Figure 1:**
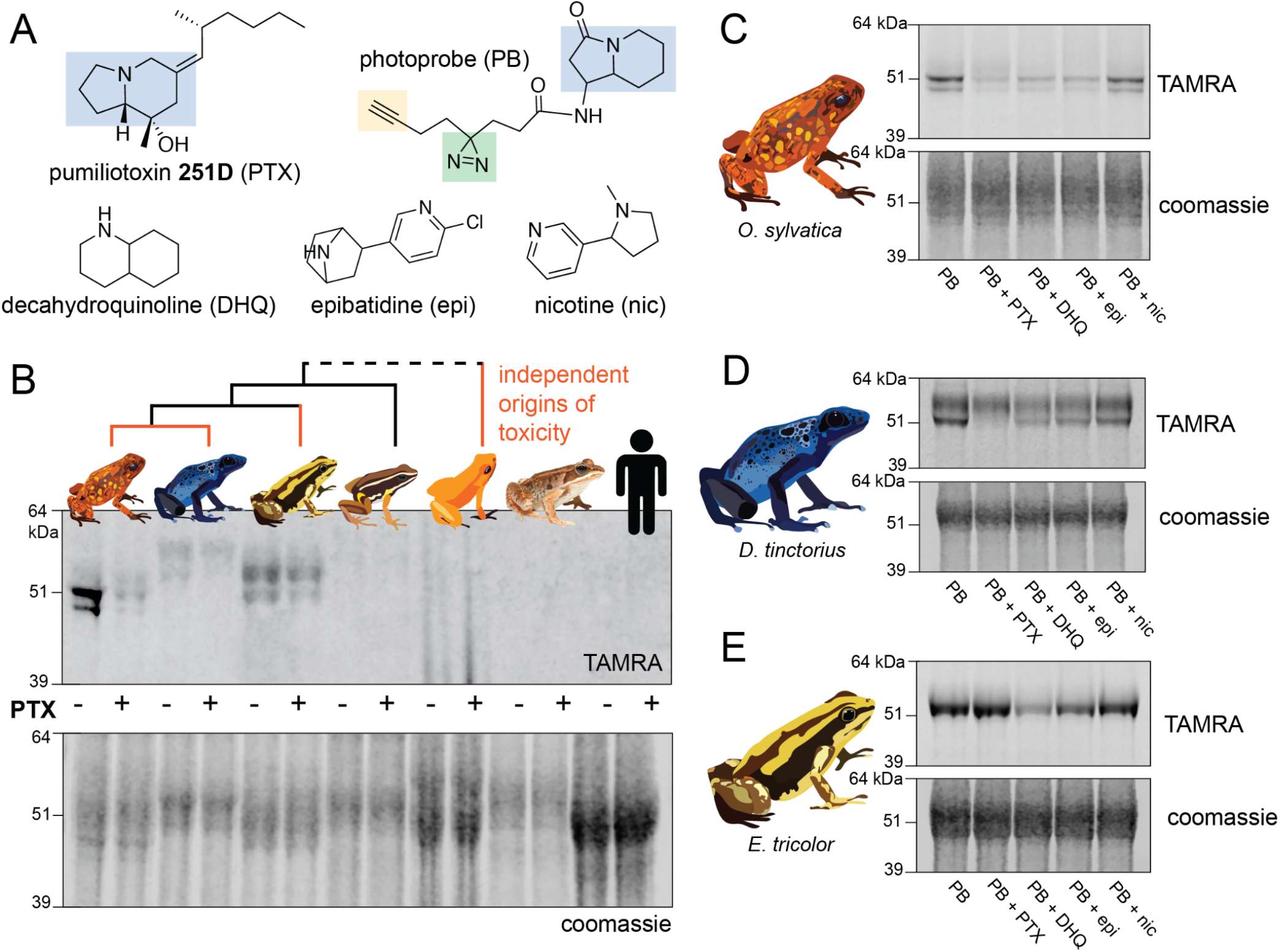
Alkaloid-like photocrosslinking probes show binding and competition in poison frog plasma. **(A)** Structures of alkaloid-like photocrosslinking probe and alkaloids tested, with the functional group in blue, the diazirine group in green, and the terminal alkyne in yellow. **(B)** Plasma from different species (*Oophaga sylvatica, Dendrobates tinctorius, Epipedobates tricolor, Allobates femoralis, Rhinella marina*, and humans, from left to right) show different plasma photoprobe binding activity and competition. Orange lines on phylogeny indicate independent evolutionary origins of chemical defense in Dendrobatidae and Mantellidae. **(C)** *Oophaga sylvatica* plasma shows crosslinking, and competition with pumiliotoxin (PTX), decahydroquinoline (DHQ), and epibatidine (epi), but not nicotine (nic). **(D)** *Dendrobates tinctorius* plasma shows crosslinking, and competition with PTX, but not DHQ, epi, or nic. **(E)** *Epipedobates tricolor* plasma shows crosslinking, and competition with DHQ and slightly with epi, but not PTX or nic.

### Proteomic analysis identifies an alkaloid-binding globulin

To identify the alkaloid-binding protein found in the plasma assays, we performed a pull-down paired with gel-punch proteomics on three conditions: no photoprobe (negative control), photoprobe only (positive control), and photoprobe with PTX competitor (Figure 2A). A biotin handle, instead of the fluorophore used above, was chemically added to the photoprobe for the enrichment of proteins using streptavidin beads. We used an untargeted proteomics approach to quantify and compare these enriched fractions using a proteome reference created from the *O. sylvatica* genome. On average, 3876 unique peptides were found per sample, mapping to 433 *O. sylvatica* proteins (Figure 2B). The most highly abundant protein in the photoprobe condition had an average number of peptide spectral counts of 1224.5 and was competed off in the photoprobe with PTX condition by 64% (Figure 2B), resembling background levels (Figure 2C). This protein is annotated as serine protease inhibitor A1 (serpin-A1), which encodes for the protein alpha-1-antitrypsin (A1AT). As our subsequent experiments demonstrate this protein functions as an alkaloid binding and sequestration protein, we refer to it hereafter as “alkaloid-binding globulin” (ABG). Mapping the 72 unique peptides onto the protein sequence of ABG showed full coverage across the protein, excluding the signal peptide, which in other serpins is cleaved during secretion (Figure S1D). We conclude that ABG functions as a major alkaloid binding protein in poison frog plasma.

**Figure 2:**
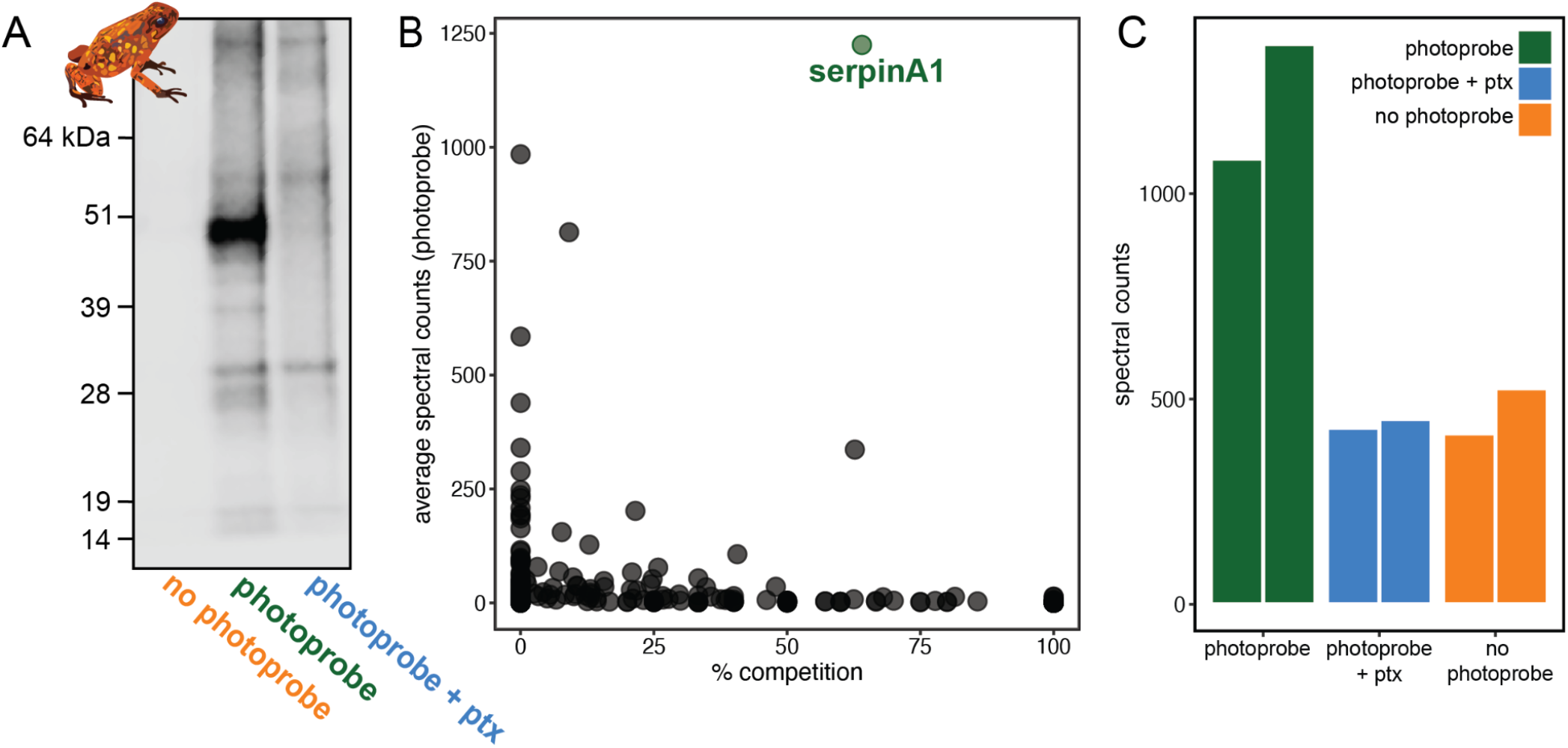
Proteomics identifies serpinA1 as the main pumiliotoxin binding protein in *Oophaga sylvatica* plasma. **(A)** Streptavidin blot of the proteins pulled down from *O. sylvatica* plasma across the three conditions: no photoprobe, photoprobe, and photoprobe plus competitor pumiliotoxin (PTX). **(B)** Quantitative proteomics output in terms of percent competition defined as 100% - average spectral counts in the photoprobe + ptx condition divided by average spectral counts in the photoprobe only condition. Average was taken across two replicates. **(C)** The number of spectral counts across conditions for the serpinA1 protein, each replicate is shown individually.

### Structural predictions of ABG show binding pocket similarities to mammalian hormone carriers

The identification of ABG as the principal alkaloid binding protein in plasma was unexpected as plasma binding of small molecules is commonly mediated by albumin. Nevertheless, in mammals, members of the serpinA family function as carriers of lipophilic hormones, providing plausibility to the hypothesis that frog serpin family members may also bind small molecules. Therefore, we sought further structural insights into ABG using protein structure predictions and molecular docking simulations to examine if this protein has a predicted binding pocket for small molecules. Using AlphaFold to predict the structure of the full protein sequence without the signal peptide resulted in a high confidence structure (Figure 3A). We then compared it to the structures of serpinA6/corticosteroid-binding globulin (CBG, Figure 3B) [39], biliverdin-binding serpin (BBS, Figure 3C) [40], serpinA1/alpha-1-antitrypsin (A1AT, Figure S2A) [41], and serpinA7/thyroxine binding globulin (TBG, Figure S2B) [42]. The AlphaFold prediction for *O. sylvatica* ABG (*Os*ABG) demonstrated a conserved structural element of three alpha helices backed by a set of beta sheets, which is the small molecule binding pocket in CBG, BBS, and TBG (black boxes, Figure 3A-C, Figure S2B), and also exists in the non-small molecule binding A1AT (black box, Figure S2A). When a molecular docking simulation was run with the whole *Os*ABG protein as the search space and PTX as the ligand, the highest affinity binding site was in the same binding pocket defined by this structural motif (Figure 3D). Although the overall structural components of the binding pockets show similarities across *Os*ABG, CBG, BBS, and TBG, the individual amino acids coordinating the small molecule binding differ across proteins (Figure 3D-F, Figure S2D-E). These results offer a structural explanation for PTX binding by ABG and highlight the homology between ABG and other small molecule binding globulins.

**Figure 3:**
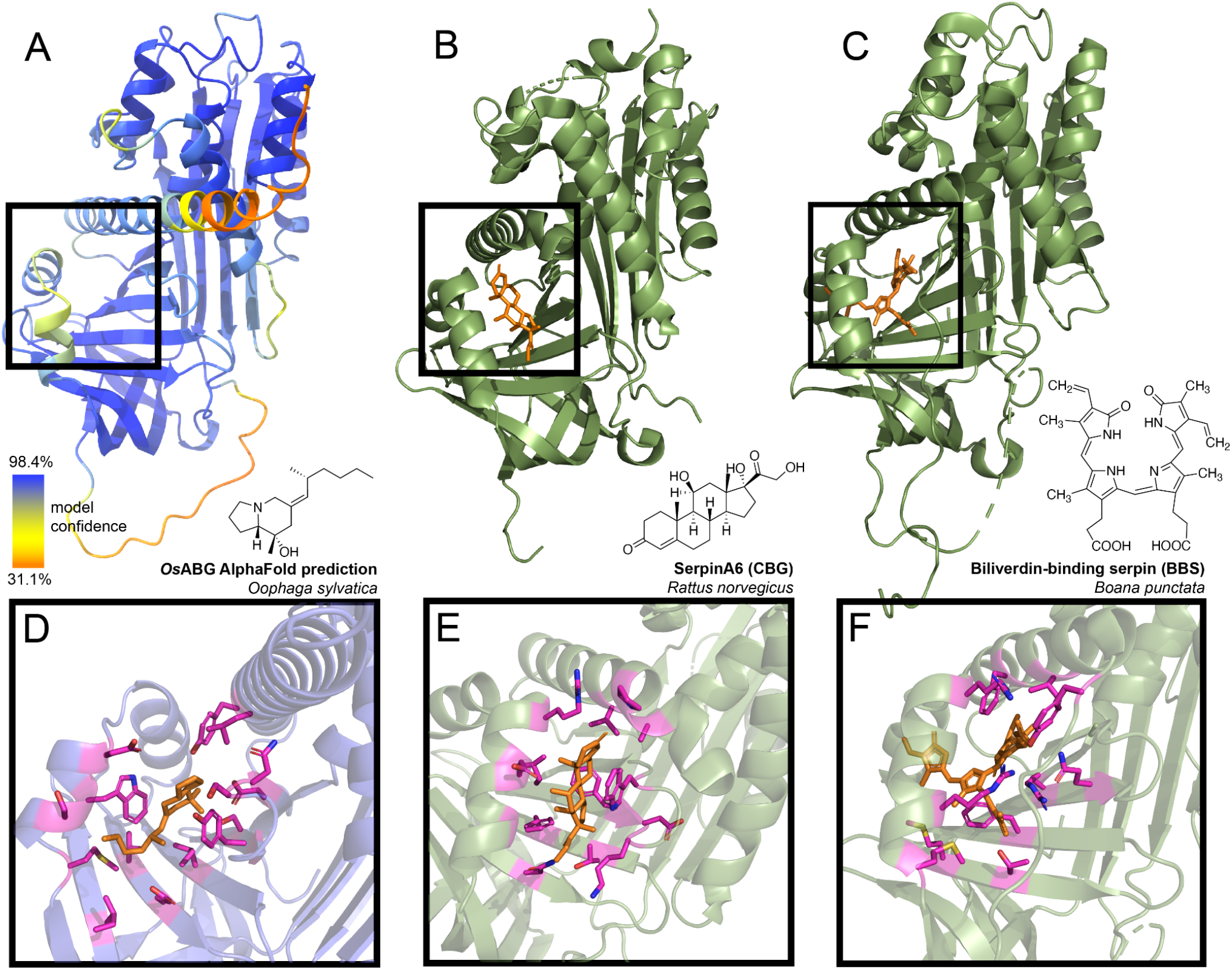
Predicted ABG structure and binding pocket resembles that of other small molecule binding serpins. **(A)** AlphaFold structure predicted with the protein sequence of the *Oophaga sylvatica* ABG, with color representing model confidence and predicted binding pocket based on molecular docking simulation indicated with a black box. **(B)** Crystal structure for rat SerpinA6/corticosteroid-binding globulin, with the cortisol molecule shown in orange (PDB# 2V95). **(C)** Crystal structure for tree frog *Boana punctata* biliverdin-binding serpin (BBS), with biliverdin shown in orange (PDB# 7RBW). **(D)** Close-up of predicted binding pocket of PTX in *O. sylvatica* ABG, with residues proximal to PTX highlighted in magenta. The structure of PTX is indicated on the top right. **(E)** Close-up of cortisol binding in CBG (PDB# 2V95), with proximal residues highlighted in magenta. Cortisol structure is displayed on the top right. **(F)** Close-up of biliverdin binding in BBS (PDB# 7RBW), with some proximal residues highlighted in magenta. Biliverdin structure is shown on the top right.

### Recombinant expression recapitulates binding activity of different ABG proteins

To confirm ABG binding activity *in vitro* and compare across different species, we recombinantly expressed and purified *O. sylvatica* ABG (*Os*ABG), and its closest homolog from the *D. tinctorius* and *E. tricolor* transcriptomes (*Dt*ABG and *Et*ABG, respectively). The resulting purified protein doublet (Figure S4B) is likely due to differences in post-translational glycosylation, as *Os*ABG has two predicted N-glycosylation sites [43]. As expected, purified *Os*ABG recapitulated the binding and competition seen with the plasma, where the photoprobe was most fully competed by the presence of PTX, and also competed off by DHQ and epibatidine, but not nicotine (Figure 4A). The competition activity with the purified protein was noticeable at a ratio of one to one photoprobe to PTX (Figure S1B). Purified *Dt*ABG and *Et*ABG required higher concentrations of protein to see a signal and showed much weaker photoprobe binding, which was competed off by the presence of PTX and DHQ in the case of *Dt*ABG (Figure 4B), and DHQ and epibatidine in the case of *Et*ABG (Figure 4C). Together these results confirm the plasma findings that ABG is a multi-alkaloid binding protein with different specificities and affinities across poison frog species, and that *Os*ABG alone is sufficient to recapitulate the crosslinking activity observed in the plasma.

**Figure 4:**
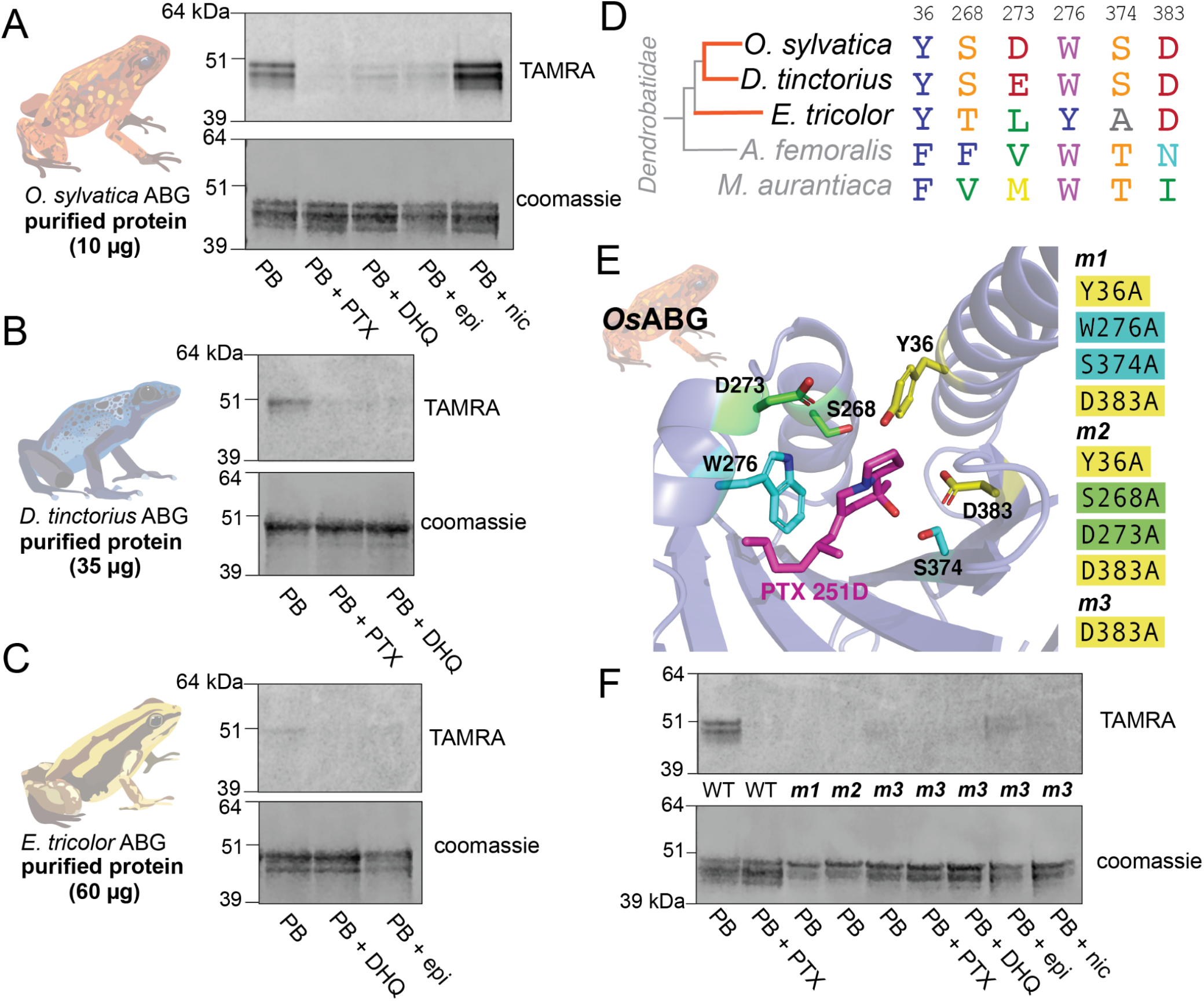
Recombinant expression and binding pocket mutants confirm plasma binding activity and binding pocket predictions. **(A)** Photoprobe crosslinking and competition with different compounds of 10 μg recombinantly expressed and purified OsABG recapitulates the binding activity seen in the plasma (Figure 1C). **(B)** Photoprobe crosslinking with 35 μg recombinantly expressed *Dendrobates tinctorius* ABG shows crosslinking, and competition with PTX and DHQ. **(C)** Photoprobe crosslinking with 60 μg recombinantly expressed *Epipedobates tricolor* ABG shows crosslinking, and competition with DHQ and epibatidine. **(D)** Alignment of protein sequence of proteins homologous to *Os*ABG across species shows conservation of certain amino acids. Coloring of amino acids is based on the RasMol “amino” coloring scheme, which highlights amino acid properties. **(E)** Potential binding residues were identified from the molecular docking simulation. Three different mutants were made based on specific amino acids in the binding pocket, with either a combination of four different alanine substitutions (m1 - yellow and teal residues, and m2 - yellow and green residues) or a single alanine substitution at D383 (m3). PTX is shown in magenta. Oxygen atoms on the molecules are highlighted in red, nitrogen in blue. **(F)** Binding pocket mutants lose binding activity of the photoprobe in the original reaction conditions.

Given the predicted binding pocket from the molecular docking simulations, and the differences in binding activity of the ABG proteins in different poison frog species, we used a sequence (Figure 4D) and predicted structure (Figure 4E) informed approach to mutate residues that might coordinate alkaloid binding in the hypothesized pocket. Using the molecular docking simulation, we identified 6 residues with proximity to the docked PTX molecule that might have important binding activity: Y36, S268, D273, W276, S374, D383 (Figure 4E). Mutating different sets of these binding residues in the *Os*ABG sequence led to a disruption of binding and competition. The combined mutations of Y36A, W276A, S374A, D383A and Y36A, S268A, D273A, D383A disrupted binding to the photoprobe completely (Figure 4E). The single point mutation of D383A weakened photoprobe binding significantly, to the point of being nearly undetectable (Figure 4E). These results demonstrate that mutating residues in the binding pocket identified through molecular docking disrupts binding activity of *Os*ABG, providing biochemical evidence that the structurally predicted binding pocket of ABG indeed is the relevant binding site for PTX. Furthermore, we have identified a set of residues that are necessary for PTX binding with high affinity, showing that the plasma binding activity is coordinated by specific amino acids in *Os*ABG.

### OsABG sequesters free PTX in solution with high affinity

Previous work has described the binding affinities of small molecule binding serpins and their important role in regulating the pool of free versus bound ligands in circulation [44–47]. We hypothesized that *Os*ABG might play a similar role for alkaloids in the poison frog plasma. To test this, we examined both the binding affinity of *Os*ABG for PTX and its ability to regulate the pool of bioavailable alkaloids in solution. Using microscale thermophoresis (MST) we found a dissociation constant (K_D_) of 1.58 μM (Figure 5A). To test the ability of *Os*ABG to sequester alkaloids *in vitro*, we used a 3 kDa molecular weight cutoff centrifuge filter to separate the “bound” and “free” PTX (Figure 5B), which we then quantified by liquid chromatography-mass spectrometry. We found that in the presence of *Os*ABG, the amount of “free” PTX is dramatically reduced, while that of nicotine is not (Figure 5C). These results show that *Os*ABG is able to bind PTX in solution with high affinity, and therefore may regulate the amount of free PTX in solution. This regulation of bioavailable pools of PTX in circulation may have downstream consequences on sequestration, transcription, and signaling throughout the organism.

**Figure 5:**
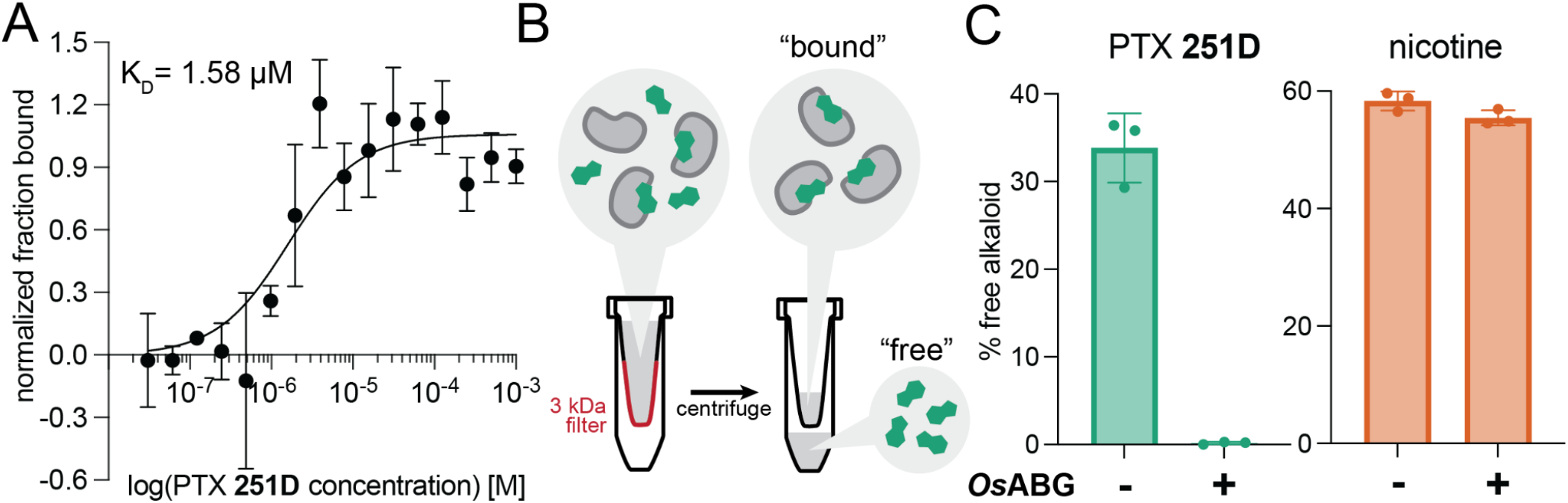
OsABG sequesters free PTX in solution. **(A)** Microscale thermophoresis (MST) of labeled *Os*ABG with PTX finds a dissociation constant (Kd) of 1.58 uM. **(B)** A 3 kDA molecular weight cut off (MWCO) centrifuge filter was used to separate “free” versus “bound” alkaloids in solutions with and without *Os*ABG present, to later be quantified with LC-MS. **(C)** The percent of “free” PTX **251D** (green) dropped when *Os*ABG was present, however the amount of “free” nicotine” remained unchanged by the presence of *Os*ABG.

### OsABG is highly expressed in wild frogs and binds ecological toxins

We next sought to better understand the functional role of OsABG in a context relevant to the ecology and physiology of poison frogs. *Oophaga sylvatica* frogs were collected across three different locations in Ecuador (Figure 6A). Tissue RNA sequencing revealed that *Os*ABG mRNA is expressed very highly in the liver compared to other tissues (Figure 6B). However a reanalysis of previously published proteomics data shows that OsABG is detectable in the liver, skin, and gut, consistent with movement through the plasma (Figure 6C). Hierarchical clustering of all unique serpinA genes in the genome shows that OsABG is most closely related to two other serpinA1 genes, Os4677 and Os4682 (Figure 6D). The liver expression of OsABG is higher than all other serpinA genes, and is higher than the expression of albumin in the liver (Figure 6D). Field-collected *O. sylvatica* frog skin contains alkaloids from many different classes, with 33% of the summed alkaloid load being histrionicotoxins, followed by 22% in 5,8-indolizidines, 15% in 3,5-indolizidines, 13% in 5,6,8-indolizidines, and 10% in decahydroquinolines (Figure 6E). Further crosslinking experiments with purified *Os*ABG found that it also binds a histrionicotoxin-like base ring structure (HTX), indolizidine (indol), and shows slight competition by a toxin mixture created from wild frog skin extracts (Figure 6F). Together, these data characterize the expression profile and distribution of OsABG and show that it is capable of binding other alkaloid classes that are found in wild frogs.

**Figure 6:**
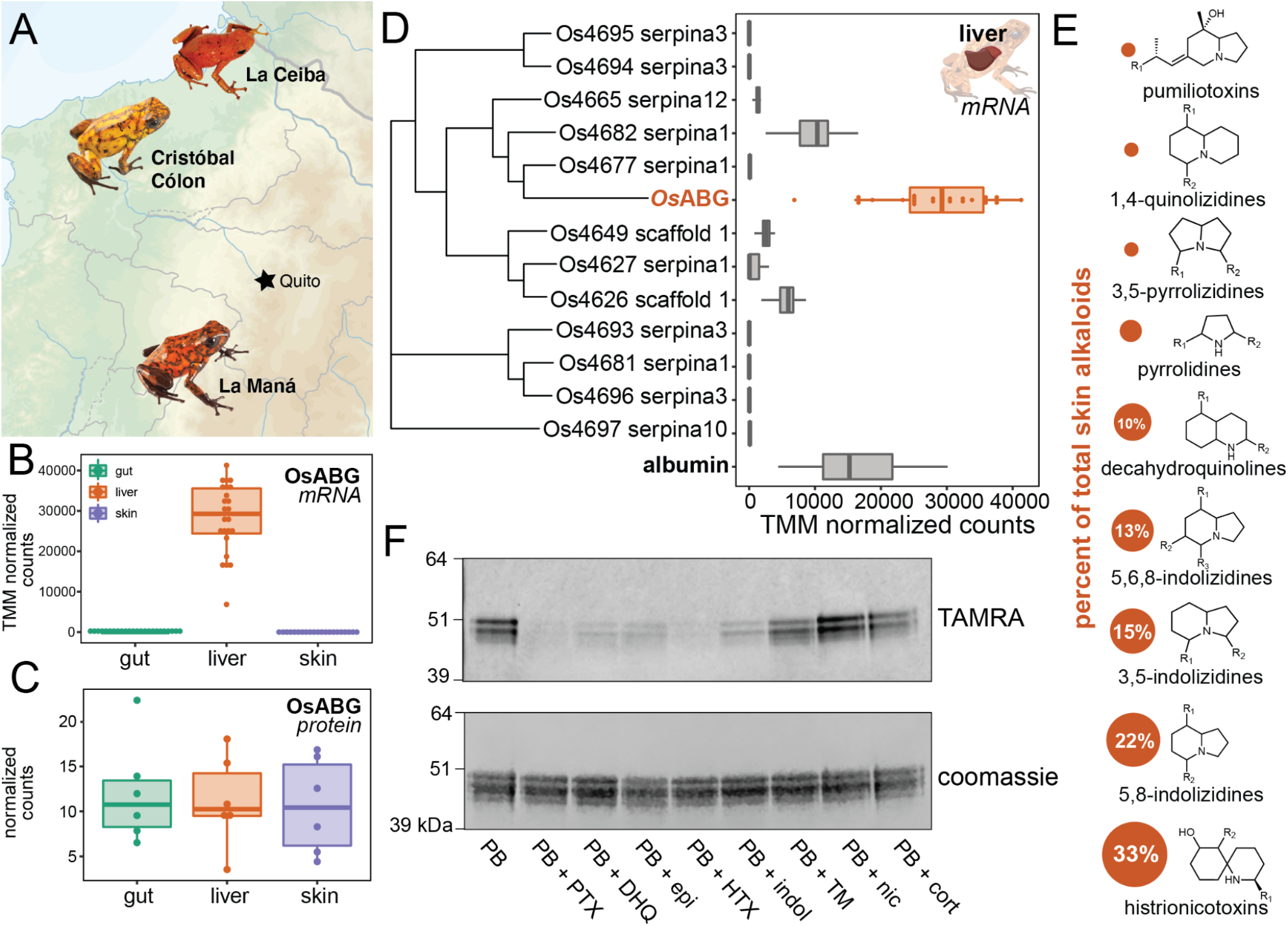
OsABG is expressed in the liver and binds ecologically relevant alkaloids. **(A)** Wild *Oophaga sylvatica* were collected across three locations in Ecuador, n=10 per location. **(B)** Comparison of mRNA expression levels across tissues found high expression in the liver, and low to no expression in the skin and gut. **(C)** Reanalysis of proteomics from Caty et al., 2022 found that OsABG protein is present in the gut, liver, and skin. **(D)** The liver expression level of OsABG was higher than that of other members of the serpinA family found in the genome, and of albumin. **(E)** Dorsal skin alkaloids fell into 9 different classes, with the size of the circle representing the averaged percent of total skin alkaloid load. **(F)** Photoprobe binding was competed by PTX, DHQ, epi, a histrionicotoxin-like compound (HTX), and indolizidine ring without R groups (indol), and slightly by a mixture of skin toxins from the wild specimens (TM). Photoprobe binding was not competed by nicotine or cortisol (cort).

## 3. DISCUSSION

Alkaloid-binding globulin (ABG) represents a new small molecule binding functionality for a member of the serpin family, with a structurally conserved binding pocket similar to mammalian hormone carriers and biliverdin-binding serpin. The broader implication of this finding is that it provides evidence for convergent evolution of serpin proteins for the binding and transport of small molecules across taxa and for different physiological requirements. Most serpin proteins are known for the anti-protease inhibitory activity, however there are members of the serpin superfamily that have been instead characterized as non-inhibitory small molecule binding globulins. In tree frogs, biliverdin-binding serpin (BBS) has been identified as the primary protein that binds biliverdin [48], the heme metabolite that is responsible for green coloration in the lineage, although it remains unclear whether BBS is also responsible for the transport of biliverdin across tissues. In mammals, cortisol-binding globulin (CBG, serpinA6) and thyroxine-binding globulin (TBG, serpinA7) play important roles in the transport and regulation of plasma hormone concentrations [49,50]. However, serpins are not the only carriers of lipophilic small molecules in mammalian plasma, with many other hormones and vitamins moving through plasma on albumin [31,32], Vitamin-D binding protein [51,52], and sex-hormone binding globulin [53,54], which have no homology to serpin proteins. That CBG and TBG have evolved small molecule transport from a protease inhibitor background [55] is surprising, and the discovery of ABG shows that this activity may have evolved either independently or from hormone transport in poison frogs for a vastly different purpose. BBS, CBG, and TBG all bind their respective ligands in a similar structural pocket [40], however the identity of the residues that coordinate binding vary across the proteins. Mutations of key binding site residues in *Os*ABG were able to disrupt binding, confirming the binding pocket predictions and the functional homology to BBS, CBG, and TBG. A more detailed mutational scanning would be necessary to fully understand which binding pocket residues coordinate the binding of different alkaloids. It would be interesting to expand binding experiments to more classes of alkaloids, although we are limited by the commercial availability of many of these compounds. The similarities between ABG and mammalian hormone carriers raises the possibility that alkaloids may also act as signaling molecules, which would require further characterization of other alkaloid targets in poison frogs and more mechanistic information about ABG alkaloid transport. The identification of this novel alkaloid carrier opens up new avenues of research in the evolution of small molecule transporters and their relevant physiological functions, especially in amphibians. The data presented suggest that OsABG is using the binding site already present in serpin family proteins, and has encoded remarkable specificity for certain alkaloids within this site.

Alkaloid Binding Globulin (ABG) has different binding specificity across poison frog species and independent evolutionary origins of acquired chemical defense, suggesting that there exist conserved mechanisms of alkaloid sequestration across the poison frog clade. Of the species tested, the photoprobe showed binding activity only in dendrobatid species that can acquire alkaloid chemical defenses from their diet, namely *O. sylvatica, D. tinctorius*, and *E. tricolor*, which represent two independent origins of chemical defense within the poison frog lineage [17]. This suggests that ABG has evolved in dendrobatid frogs that are capable of acquired chemical defense, as binding was not seen in the dendrobatid without chemical defenses in nature (*A. femoralis*), the Malagasy poison frog (*M. aurantiaca*), cane toads, or human plasma. Malagasy poison frogs represent a convergent evolution of many of the notable phenotypes in the dendrobatid lineage, and have been found to contain pumiliotoxins and other dendrobatid alkaloids [56–58]. The lack of binding activity in the plasma may indicate that they have evolved different molecular mechanisms for alkaloid transport and autoresistance. Using competition of the photoprobe by presence of excess alkaloid as a proxy for binding activity, we found differences in the competition activity of different alkaloids across species. This different plasma binding suggests that *O. sylvatica* has a more promiscuous alkaloid binding pocket, which may be related to the high diversity of alkaloids found in wild *O. sylvatica* frogs [27,59,60]. Future characterization of plasma binding activity with more species and alkaloid compounds would be critical to fully trace the evolution of ABG binding across the poison frog lineage. The identification of these three ABG proteins opens up the possibility of further discoveries of ABGs in other toxic species, and a more comprehensive understanding of the biochemical parameters involved in binding structurally different ligands. Overall, the diversity in plasma binding seen across phylogenetically close species may reflect the diversity in environmental pressures that have led to species-specific adaptations at the molecular scale.

*Os*ABG is expressed at high levels and binds additional alkaloids found in high abundance on the skin of field-collected *O. sylvatica*, indicating its physiological and ecological relevance for this poison frog species. *Os*ABG is produced in the liver as is the case with most members of the serpin family in humans [61], cows [62], rats [63], baboons [64], and macaques [62]. The liver expression level of *Os*ABG was much higher than other serpinA proteins and albumin, which in most vertebrates is the most abundant plasma protein [32,65,66], further highlighting the importance of this protein for poison frog physiology. Taken together, this supports a model where *Os*ABG is produced in the liver, binds alkaloids present in the blood or recently absorbed by the intestines, and then may transport alkaloids to the skin for bioaccumulation. The micromolar K_D_ value we identified for *Os*ABG with PTX is higher than the previously reported value for CBG [44,46], and TBG [67], as well as that reported for the “toxin sponge” protein saxiphilin [68], which are all in the nanomolar range. The lower affinity of *Os*ABG provides further support to the hypothesis that *Os*ABG may be acting as a transporter protein to other tissues, and would be in line with the hypothesis that there may be other mechanisms involved in autoresistance to circulating alkaloids not bound by protein [69]. It is also possible that *Os*ABG has affinities in the nanomolar range for other ligands which were not tested in this study. Previous work with CBG and TBG in mammals show that they bind 70-90% of their respective ligands in the plasma, thus regulating the concentration of free hormone in the blood [50,70]. Furthermore, CBG and TBG have transport activity, as the release of the ligand is induced by proteolytic cleavage of the reactive center loop which creates a conformational change of the protein [49,71]. ABG may be acting in a similar way to mediate the bioaccumulation of dietary alkaloids onto the skin. The reduction of “free” PTX when *Os*ABG is present *in vitro* further supports this, given that *Os*ABG would be regulating the pool of unbound alkaloids available in the plasma. Further characterization using the respective protease from *O. sylvatica* would be necessary to validate the alkaloid release through proteolytic cleavage of the reactive center loop as well as genetic knockout frogs to fully characterize the organismal role of *Os*ABG. Given these findings we conclude that ABG may play this important transport role in poison frog physiology and chemical defense, which greatly broadens our understanding of this previously elusive mechanism in poison frogs. Furthermore, given its striking similarities with mammalian hormone carriers and biliverdin binding serpin, our findings open up additional questions regarding the functional convergence of serpins as small molecule transporters in distantly related taxa.

### Summary

This study presents the first evidence of an alkaloid binding protein in the poison frog plasma with a suggested role as a transporter molecule. We found a novel alkaloid binding function for a member of the serpin family, contributing to a mounting body of evidence suggesting that the small molecule binding activity of certain serpins has evolved multiple times from their protease inhibitor precursors with a structurally conserved binding pocket. Our alkaloid comparisons with *Os*ABG show that this serpin binding pocket has been fine-tuned for specificity for certain small molecule substrates, and may play an important ecological and physiological role in the evolution of chemical defense in poison frogs. The discovery of a serpin that can bind a number of different ligands sets up the possibility of future efforts to understand and engineer ABG’s binding pocket for new ligands once an in-depth understanding of the binding specificity is achieved.

## 4. MATERIALS AND METHODS

### Animal usage

All animal procedures were approved by the Institutional Animal Care and Use Committee at Stanford (protocol #34153). Topical benzocaine was used for anesthesia prior to the euthanasia of all animals. Laboratory-bred animals were purchased from Understory Enterprises (Ontario, Canada) or Josh’s Frogs (Michigan, USA) depending on the species. Animals were either euthanized for plasma collection upon arrival, or housed in 18^3^ inch glass terraria, and fed a diet of non-toxic *Drosophila melanogaster* until euthanasia. Plasma and tissues from a total of 62 animals were used for this study, consisting of 32 lab-bred animals and 30 field-collected animals that are described below. Sample size of field-collected animals was determined based on variability seen in previous studies, and for laboratory experiments based on experimental needs in terms of volume of plasma.

### Plasma collections

Lab-bred, and therefore non-toxic, poison frogs were anesthetized with topical application of 10% benzocaine on the ventral skin, and euthanized with cervical translocation. Blood was collected directly from the cervical cut using a heparinized capillary tube (22-362-566, Fisher Scientific, Waltham, MA) and deposited into a lithium heparin coated microvette tube (20.1282.100, Sarstedt, Nümbrecht, Germany). Blood was spun down 10 minutes at 5000 rpm at 4°C on a benchtop centrifuge, and the top layer which contains the plasma was removed and pipetted into a microcentrifuge tube. This was stored at −80°C until it was used for experiments.

### UV crosslinking and competition using alkaloid-like photoprobe

Photocrosslinking methods follow methods outlined in Kim *et al*., 2020 [72]. Plasma or purified protein was thawed on ice. The total reaction volume was 50 μL and all experiments were performed in a clear 96 well plate. For plasma, 5 μL of undiluted plasma was mixed with 40 μL of PBS for each reaction. For purified *O. sylvatica* protein, 10 μg of protein was diluted into PBS per reaction to a volume of 45 μL. For *E. tricolor* and *D. tinctorius* protein, 60 μg and 35 μg were used, respectively; a higher amount of protein was used because no photocrosslinking was detected at 10 μg for these proteins. To this, either 2.5 μL of DMSO was added as vehicle control, or 2.5 μL of competitor compound dissolved in DMSO was added at a final concentration of 100 μM for plasma competition experiments unless indicated otherwise, or 1 mM for purified protein experiments unless indicated otherwise. The competitor compounds were: custom synthesized pumiliotoxin **251D** (PTX, PepTech, Burlington, MA), decahydroquinoline (DHQ, 125741, Sigma-Aldrich, St. Louis, MO), epibatidine (epi, E1145, Sigma-Aldrich), a histrionicotoxin-like compound (HTX, ENAH2C55884A-50MG, Sigma-Aldrich), indolizidine (indol, ATE24584802-100MG, Sigma-Aldrich), nicotine (nic, N3876-100ML, Sigma-Aldrich), cortisol (cort, H0888-1G, Sigma-Aldrich). The “toxin mixture” (TM) used as a competitor in Figure 6 was made by taking 20μL of each of the skin alkaloid extracts from wild frogs described below, evaporating it under gentle nitrogen gas flow, and resuspending in 100μL of DMSO. This was followed by addition of 2.5 μL of photoprobe (Z2866906198, Enamine, Kyiv, Ukraine) dissolved in DMSO on ice, for a final photoprobe concentration of 5 μM in plasma experiments and 100 μM in purified protein experiments. This was incubated on ice for 10 minutes, and then UV crosslinked (Stratalinker UV 1800 Crosslinker, Stratagene, La Jolla, CA) for 5 minutes on ice. TAMRA visualization of crosslinked proteins was done by adding 3 μl TBTA (stock solution: 1.7 mM in 4:1 v/v DMSO:tert-Butanol; H66485-03, Fisher), 1 μl Copper (II) Sulfate (stock solution: 50 mM in water; BP346-500, Fisher), 1 μl Tris (2-carboxyethyl) phosphine hydrochloride (freshly prepared, stock solution: 50 mM; J60316-06, Fisher), and 1 μl TAMRA-N3 (stock solution: 1.25 mM in DMSO; T10182, Fisher), incubating at room temperature for 1 hour, and quenching the reaction by boiling with 4x SDS loading buffer. This was run on a Nupage 4-12% Bis-Tris protein gel (NP0323BOX, Invitrogen, Waltham, MA) and the in-gel fluorescence of the gel was visualized using a LI-COR Odyssey imaging system (LI-COR Biosciences, Lincoln, Nebraska) at 600 nm for an exposure time of 30 seconds. After imaging the TAMRA signal, the same gel was coomassie stained (InstantBlue, ISB1L, Abcam, Cambridge, UK) and visualized the same way at 700 nm.

For proteomic identification of competed proteins, plasma samples were pooled from five different individuals and were crosslinked with either no photoprobe and equivalent amounts of DMSO, 5 μM photoprobe and DMSO, or 5 μM photoprobe and 100 μM PTX as described above. Each condition was set up as 24 individual reactions and pooled after crosslinking. To attach a biotin handle, 3 μl TBTA, 1 ul CuSO4, 1 μl TCEP, and 1.14 μl Biotin-N3 (stock solution: 9.67 mM in DMSO; 1265, Click Chemistry Tools, Scottsdale, AZ), were added for each reaction and this was incubated at room temperature for 1 hour, rotating. After incubation, each condition was run through a 3kDa MWCO centrifuge filter twice (UFC800324, Amicon, Millipore-Sigma, Burlington, MA) to dilute excess Biotin-N3 until reaching a 900x dilution. Pulldown of biotinylated photoprobe-protein complexes was achieved with a magnetic bead strep pulldown following the protocol outlined in Wei *et al*., 2021[73]. The pre and post-pulldown samples were run on a gel for a streptavidin blot (Figure 2A) and silver stain. After verifying pulldown efficacy, samples were run on SDS-PAGE gels in two replicates (one lane each) for each condition. Gels were fixed in 50:50 water:MeOH with 10% Acetic Acid for 1-2 hours. For the first replicate, the gel was run for a short period, and an approximately one centimeter squared portion containing the whole lane for each condition was excised and fixed. For the second replicate, the gel was run completely and the proteins between 39 kDa and 64 kDa were excised and fixed using the ladder as a size reference. Sections were chopped into 1 mm pieces under sterile conditions and stored at 4°C in 100μL of water with 1% acetic acid until processed for proteomics.

### Proteomic identification of pulled down proteins across conditions

For proteomics analyses, SDS-PAGE gel slices approximately 1 cm in length were prepared for proteolytic digestion. Each fixed gel slice was diced into 1 mm cubes under sterile conditions, and then rinsed with 50 mM ammonium bicarbonate to remove residual acidification from the fixing process. Following rinsing, the gels were incubated in 80% acetonitrile in water for five minutes; the solvent was removed and then the gel pieces were incubated with 10 mM DTT dissolved in water at room temperature for 20 minutes. Following reduction, alkylation was performed using 30 mM acrylamide for 30 minutes at room temperature to cap free reduced cysteines. Proteolysis was performed using trypsin/lysC (Promega, Madison, WI) in 50 mM ammonium bicarbonate overnight at 37°C. Resulting samples were spun to pellet gel fragments prior to extraction of the peptides present in the supernatant. The resulting peptides were dried by speed vac before dissolution in a reconstitution buffer (2% acetonitrile with 0.1% formic acid), with an estimated 1 μg on-column used for subsequent LC-MS/MS analysis.

The liquid chromatography mass spectrometry experiment was performed using an Orbitrap Eclipse Tribrid mass spectrometer RRID:022212 (Thermo Scientific, San Jose, CA) with liquid chromatography using an Acquity M-Class UPLC (Waters Corporation, Milford, MA). A flow rate of 300 nL/min was used, where mobile phase A was 0.2% formic acid in water and mobile phase B was 0.2% formic acid in acetonitrile. Analytical columns were prepared in-house with an I.D. of 100 microns pulled to a nanospray emitter using a P2000 laser puller (Sutter Instrument, Novato, CA). The column was packed using C18 reprosil Pur 1.8 micron stationary phase (Dr. Maisch) to an approximate length of 25 cm. Peptides were directly injected onto the analytical column using a 80 minute gradient (2%–45% B, followed by a high-B wash). The mass spectrometer was operated in a data dependent fashion using CID fragmentation in the ion trap for MS/MS spectra generation.

For data analysis, the .RAW data files were processed using Byonic v4.1.5 (Protein Metrics, Cupertino, CA) to identify peptides and infer proteins based on a proteomic reference created with the *O. sylvatica* genome annotation. Proteolysis with Trypsin/LysC was assumed to be specific with up to 2 missed proteolytic cleavages. Precursor mass accuracies were held within 12 ppm, and 0.4 Da for MS/MS fragments in the ion trap. Cysteine modified with propionamide were set as fixed modifications in the search, and other common modifications (e.g. oxidation of methionine) were also included. Proteins were held to a false discovery rate of 1%, using standard reverse-decoy technique [74].

### Identification of ABG proteins in different species and sequence confirmation

To identify potential ABG proteins in other species, we used the *Os*ABG protein sequence identified in the proteomics as the query and searched against blast databases created from the *Allobates femoralis* genome, and *Epipedobates tricolor*, *Dendrobates tinctorius*, and *Mantella aurantiaca* transcriptomes. The top hit from each blast search was used as the most probable ABG gene from those species. To ensure that the sequences did not contain sequencing or alignment errors, the gene from *O. sylvatica, D. tinctorius, and E. tricolor* was amplified using PCR and sequence confirmed with sanger sequencing. Total RNA was extracted from flash frozen liver tissue from three lab-bred, non-toxic, individuals from each species using the Monarch total RNA Miniprep Kit (T2010S, New England Biolabs, Ipswich, MA) following manufacturer instructions. This was used to create cDNA using the SuperScript III First-Strand Synthesis kit (18080-400, Invitrogen), following manufacturer instructions with an oligo(dT)20 primer. This was used for a PCR using Phusion High Fidelity DNA polymerase (F-530, Thermo Scientific) and the primers and cycling conditions described below. PCRs were analyzed using a 1% agarose gel for presence of a single band, cleaned up (NucleoSpin Gel and PCR cleanup, 740609.50, Takara Bio, Shiga, Japan), and transformed into pENTR vectors using a D-TOPO kit (45-0218, Invitrogen). Plasmids containing the ABG sequences from each individual were then mini prepped (27106X4, Qiagen, Hilden, Germany) and sanger sequenced with M13F and M13R primers (Azenta Life Sciences, South San Francisco, California). Sequences were aligned using Benchling (Benchling Inc, San Francisco, California) software, with MAFFT used for DNA alignments and Clustal Omega used for protein alignments.

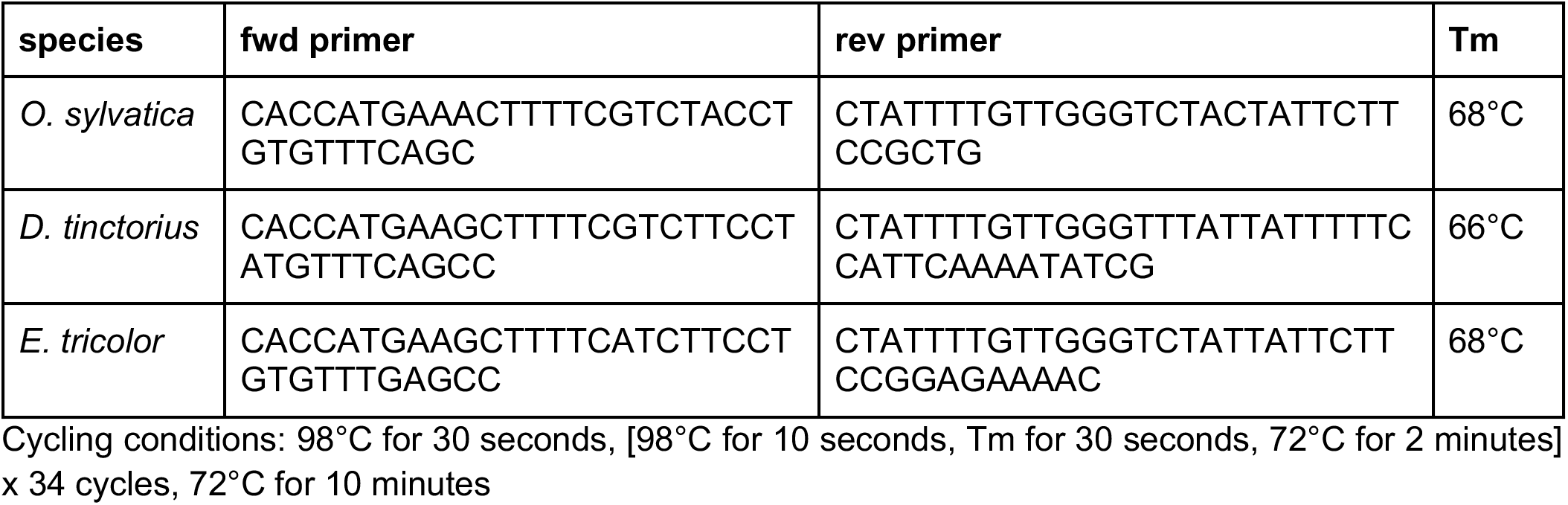

### Protein structure prediction and molecular docking analyses

The *Os*ABG protein folding was predicted using the amino acid sequence, edited for point mutations found across all three individuals used for sequence verification, and the AlphaFold google colab notebook (https://colab.research.google.com/github/deepmind/alphafold/blob/main/notebooks/AlphaFold.ipynb)[7 5]. The predicted structure is provided in the supplementary information. The default AlphaFold parameters were used. Molecular docking was performed using the UCSF Chimera software (https://www.cgl.ucsf.edu/chimera/)[76], using AutoDock Vina [77,78] with the three dimensional structure of PTX **251D** (Pubchem CID 6440480). The whole protein was used as the search space with the default search parameters (5 binding modes, exhaustiveness of search of 8, and a maximum energy difference of 3 kcal/mol). The docking result with the highest predicted affinity was used and is included in the supplementary files. Protein structures and docking were visualized using PyMol for publication quality images.

### Recombinant protein expression and binding assays

Recombinant ABG proteins were expressed by Kemp Proteins (Maryland, USA) through their custom insect cell protein expression and purification services. The reagents and vectors used are proprietary to Kemp Proteins, however the general expression and purification details are as follows. The verified protein sequences described above or the point mutations (Figure 4) were codon-optimized for SF9 insect expression, and a 10xHIS tag was added to the C-terminal end. For *Os*ABG, a 1 L expression was performed, for all other sequences (other species and mutants) a 50 mL expression was used. For the 1 L expression, a multiplicity of infection of one was used for the p1 baculovirus and the supernatant was collected after 72 hours. To this, 5 mL of Qiagen Ni-NTA resin washed and equilibrated in Buffer A (20 mM Sodium Phosphate, 300 mM NaCl, pH=7.8) was added and it was mixed overnight at 4°C. Afterwards, it was packed in a 5 mL Bio-Scale column and washed with 3 column volumes (CV) of Buffer A, followed by washing with 5% Buffer B (20 mM Sodium Phosphate, 300 mM NaCl, 500 mM Imidazole, pH=7.8) for 5 CV. Protein was eluted with a linear gradient from 5-60% over 25 CV, and 6 mL fractions were collected throughout. All of the fractions containing protein were pooled and concentrated to 1 mg/mL using an Amicon centrifugal filter with a 10 kDa MWCO, the buffer was exchanged to PBS, it was filtered through a 0.2 um filter, aliquoted, and frozen at −80°C. Protein expression and purification resulted in a clear band by western blot (Figure S4A) and a clean doublet pattern by coomassie (Figure S4B) closely resembling that seen in the plasma crosslinking results (Figure 1C) in both reduced and non-reduced conditions. For the 50mL expression of *Dt*ABG, *Et*ABG, and mutant *Os*ABG proteins, a 10% ratio of p1 virus to media was used and the supernatant was collected after 72 hours, to which 1 mL of Qiagen Ni-NTA resin washed and equilibrated in Buffer A (20 mM Sodium Phosphate, 300 mM NaCl, pH=7.4) was added. This was mixed overnight at 4°C and then packed into a 1 mL Bio-Scale column, washed with 3 CV of Buffer A, washed with 5 CV of 5% Buffer B (20 mM Sodium Phosphate, 300 mM NaCl, 500 mM imidazole, pH=7.4), and eluted with 5 CV of 50% Buffer B. Fractions containing protein were buffer exchanged into PBS, and the final concentrations were approximately 0.2 mg/mL, with varying final volumes. Protein expression and purification resulted in a clear band by western blot (Figure S4C,E,G,I,K), and a clean doublet by coomassie (Figure S4D,F,H,J,L) in both reduced and non-reduced conditions.

### Determination of dissociation constant using Microscale Thermophoresis (MST)

To determine the binding affinity of *Os*ABG for PTX, we used Microscale Thermophoresis (MST) to determine the dissociation constant (K_D_). To do this, we used the Monolith system (Nanotemper Technologies, München, Germany). Purified *Os*ABG protein was labeled using the protein labeling kit Red-NHS 2nd generation (MO-L011, Nanotemper) which dyes primary lysine residues in the protein. The kit was used following manufacturer instructions, however a 1.5x excess of dye was used instead of 3x as this was found to better achieve a degree of labeling of ~0.5. To remove aggregates during the assay, PBS-Tween was used for protein labeling and all dilutions. The labeled protein was centrifuged for 10 minutes at 20,000g on a benchtop centrifuge, and the supernatant was taken to further remove any aggregation. The concentration was measured prior to calculating and setting up dilution series. A final concentration of 10nM *Os*ABG was used, and a 16 tube 2x serial dilution series of PTX 251D was made with the highest concentration being 1000uM. The concentration of DMSO was maintained consistent across the dilution series. The labeled *Os*ABG was incubated for 10 minutes prior to loading into capillaries, and three biological replicates were pipetted and run separately. The Monolith premium capillaries (MO-K025, Nanotemper) were used, the MST power was set to Medium, and the excitation power was set to auto-detect. The three replicates were compiled and plotted together using GraphPad Prism (GraphPad Software, San Diego, California), and a dissociation model was fit to the data to calculate the K_D_. The raw data is included in the supplementary information (will be included with full submission).

### Determination of free versus bound alkaloids

Solutions with 4 uM of *Os*ABG protein, 4 uM of either PTX **251D** or nicotine, and a final volume of 100 uL were made and incubated for one hour at room temperature. This was transferred to a 3 kDa MWCO centrifugal filter (UFC500396, Amicon) and spun at max speed on a benchtop centrifuge at 4C for 45 minutes. The top and bottom fractions were brought up to 100 uL with ultrapure water and transferred to new tubes, where 300 uL of 2:1 Acetonitrile:Methanol was added, after which they were vortexed and centrifuged at max speed on a benchtop centrifuge at 4C for 10 minutes. The supernatant was transferred to autosampler vials for quantitation of the amount of alkaloid in each fraction with mass spectrometry. Each condition was run in triplicate. Samples were analyzed using an Agilent Quadrupole time-of-flight LC-MS instrument, with MS analysis performed by electrospray ionization (ESI) in positive mode. Metabolites were separated with an Eclipse Plus C18 column (Agilent 959961-902) with normal phase chromatography. Mobile phases were: buffer A (water with 0.1% formic acid) and buffer B (90% acetonitrile, 10% water with 0.1% formic acid). The flow rate was maintained constant at 0.7 mL/min throughout the LC protocol. The LC gradient elution was set as follows: starting at 5% B held till 0.51 minutes, linear gradient from 5 to 25% B in 1.5 minutes, linear gradient from 25 to 50% B in 23 minutes, linear gradient from 50 to 95% B in 30 seconds, 95% B held for 2 minutes, linear gradient from 95 to 5% B in 1 minute, and 5% B held for 1.5 minutes to equilibrate the column to the initial conditions. The total run time was 30 minutes and the injection volume was 10 uL. Data was analyzed using the Agilent MassHunter software; the extracted ion chromatograms for PTX were searched using the exact mass M+1 of 252.2333, and nicotine was searched using the exact mass M+1 of 163.123, with a tolerance of a symmetric +/- 100 ppm. Extracted ion chromatograms were smoothed once before automatically integrating to get the abundance values. Abundance values were used to calculate the fractions above and below the filter for each replicate, which were then plotted with GraphPad. All raw data is provided as mzXML files through DataDryad (accessing information will be added for full submission)..

### Field collections of *Oophaga sylvatica*

The frog samples used in this paper are the same as those used for the project described in Moskowitz *et al*., 2022 [79]. For each location, 10 *O. sylvatica* individuals were collected under collection permit 0013-18 IC-FAU-DNB/MA issued by the Ministerio del Ambiente de Ecuador, between the hours of 7:00-18:00 during early May to early June 2019. All individuals were euthanized the same day as collection. Prior to euthanizing, frogs were sexed, weighed, and the snout-vent length was measured. Orajel (10% benzocaine) was used as an anesthetic prior to cervical dislocation. Once euthanized, frogs were immediately dissected and the liver, intestines, and half of the dorsal skin were stored in RNAlater in cryotubes at room temperature. The other half of the dorsal skin was placed in methanol in glass tubes at room temperature. Once back in the lab, all tissues were stored at −20°C until further processing. All tissues were transported to the United States under CITES permits 19EC000036/VS, 19EC000037/VS, 19EC000038/VS.

### Alkaloid extraction, detection, and analysis

All following steps were performed under a hood. Skins were taken out of methanol with forceps and weighed. From the methanol that the skin was stored in, 1 mL was taken and syringe filtered through a 0.45μ PTFE filter (44504-NP, Thermo Scientific) into the new glass vial with a PTFE cap (60940A-2, Fisher) filled with 25 μL of 1 μg/μL (-)-Nicotine (N3876-100ML, Sigma Aldrich), for a total of 25 μg of added nicotine. Tubes were capped and vortexed, and stored at −80°C celsius for 24 hours, during which proteins and lipids should precipitate. After 24 hours, tubes were taken out of the −80°C and quickly syringe filtered through a 0.45μ PTFE filter again into a new glass vial. A 100 μL aliquot was added to an GC-MS autosampler vial, and remaining solution was stored in the original capped vial at −80°C.

GC-MS analysis was performed on a Shimadzu GCMS-QP2020 instrument with a Shimadzu 30m x 0.25 mmID SH-Rxi-5Sil MS column closely following the protocol outlined in Saporito *et al*., 2010 [80]. In brief, GC separation of alkaloids was achieved using a temperature program from 100 to 280°C at a rate of 10°C per minute with He as the carrier gas (flow rate: 1 mL/min). This was followed by a 2 minute hold and additional ramp to 320°C at a rate of 10°C/minute for column protection reasons, and no alkaloids appeared during this part of the method. Compounds were analyzed with electron impact-mass spectrometry (EI-MS). The GC-MS data files were exported as CDF files and the Global Natural Products Social Network (GNPS) was used to perform the deconvolution and library searching against the AMDIS (NIST) database to identify all compounds (https://gnps.ucsd.edu) [81]. For deconvolution (identification of peaks and abundance estimates) the default parameters were used, for the library search the precursor ion mass tolerance was set to 20000 Da and the MS/MS fragment ion tolerance to 0.5 Da. The resulting dataset was filtered to keep only compounds that matched to our spiked-in nicotine standard, alkaloids previously found in poison frogs from the Daly 2005 database [15], or compounds with the same base ring structure and R groups as the classes defined in Daly 2005. All GC-MS data as CDF files are available through the GNPS public data repository (accessing information will be added for full submission).

Once the feature table from the GNPS deconvolution was filtered to only include only poison frog alkaloids and nicotine, the abundances values were normalized by dividing by the nicotine standard and skin weight. This filtered and normalized feature table was used for all further analyses and visualizations. All steps were carried out with R version 4.0.4, and code is included in supplementary data (will be included with full submission).

### RNA extraction and library preparation

RNA extraction followed the Trizol (15596018, Thermo Fisher) RNA isolation protocol outlined in Caty *et al*. 2019 [27] according to the manufacturer’s instructions, and with sample randomization to avoid batch effects. RNA quality was measured on a Agilent Tapestation RNA screentape (Agilent, Santa Clara, CA), and quantified using a Qubit Broad Range RNA kit (Q10210, Invitrogen). In the liver and intestines, samples with RIN scores greater than 5 were kept, RNA was normalized to the same Qubit concentration, and mRNA was isolated and library prepped using the NEB Directional RNA sequencing kit (E7765L, New England Biolabs) with the PolyA purification bundle (E7490L, New England Biolabs) and 96 Unique Dual Indices (E7765L, New England Biolabs). The skin RIN scores were much lower, signaling potential RNA degradation, ribosomal degradation was instead used to isolate mRNA. Following normalization within all skin RNA samples to the same Qubit concentration, we used the Zymo RiboFree Total RNA Library Prep kit (R3003-B, Zymo Research, Irvine, CA) following manufacturer instructions. After library prep for all tissues was complete library size was quantified with the Agilent Tapestation D1000 screentape, and concentration was measured with the Qubit dsDNA high sensitivity kit (Q33231, Invitrogen). All libraries within a tissue type were pooled to equimolar amounts and sequenced on two lanes of an Illumina NovaSeq (Illumina, San Diego, CA) machine to obtain 150 bp paired-end reads.

### RNA expression analysis and identification of *O. sylvatica* serpinA genes

Analysis of RNA expression levels followed the protocol outlined by Payne *et al*., 2022 [82]. The Trim-galore! wrapper tool [83] was used to trim adapter sequences with cutadapt [84] and quality filter the reads (trim_galore --paired --phred33 --length 36 -q 30 --stringency 1 -e 0.001). All trimmed reads are available through the NCBI BioProject (accessing information will be added for full submission). *Kallisto* [85] was used to pseudoalign the reads to a reference created with the coding sequence of the annotated *O. sylvatica* genome. These abundances were combined into a matrix, and the trimmed-mean of M-values (TMM) normalized counts were used for all further analyses. Additional serpinA genes were found in the genome by searching for all genes annotated with “serpina” in the header, and by blasting the *Os*ABG protein sequence against the genome (e-value < 1e-60) and including any additional genes not annotated with “serpina.” Four sequences were removed because they were exact matches of the full gene (OopSylGTT00000004683), the N-terminal end (OopSylGTT00000004650, OopSylGTT00000004685), or the C-terminal (OopSylGTT00000004676) end sequence of another serpina gene, and therefore could be potential duplications caused by annotation or assembly errors. To create the protein tree (Figure 6D), ClustalW was used to align the sequences, a distance matrix was created using identity, and neighbor joining was used to construct the tree. The albumin gene was determined by blasting the protein sequences of *Xenopus laevis* albumin A (Uniprot #P08759), *X. laevis* albumin B (Uniprot #P14872), and the Asian toad *Bombina maxima* albumin (Uniprot #Q3T478) against the *O. sylvatica* genome. In all three cases, the top hit was the same (OopSylGTT00000003067), therefore this was assumed to be the most likely albumin candidate in the genome and was used to plot the TMM expression for comparison. All plots were created in R version 4.0.4, and all analysis and plotting code is available in the supplementary files (will be included with full submission).

## 5. ACKNOWLEDGEMENTS

The authors acknowledge that this research was conducted on the ancestral lands of the Muwekma Ohlone people at Stanford; we understand the implications of the historical and present colonialism experienced by the Ohlone people and celebrate their continued stewardship of their lands. We thank Andrea Terán Valdéz for their assistance coordinating field work and their kindness. We would also like to thank the Laboratory of Organismal Biology, the Long Lab, the Soh lab, Joel Francis, and Cheyenne Payne for helpful discussions and guidance throughout this project. Centro Jambatu researchers thank the Saint Louis Zoo for their commitment and sustained support for amphibian research.

## 7. DATA AVAILABILITY

All raw data, analysis scripts, and intermediate data analysis files will be publicly available at the time of publication.

## 9. AUTHOR CONTRIBUTIONS

AAB - Conceptualization, Methodology, Validation, Formal Analysis, Investigation, Data Curation, Writing - Original Draft, Writing - Review and Editing, Visualization

MDMG - Methodology, Validation, Formal Analysis, Investigation

AER - Methodology, Validation, Formal Analysis, Investigation

ET - Methodology, Validation, Investigation

JT - Methodology

LAC - Methodology, Resources

HTS - Methodology, Resources

JZL - Conceptualization, Methodology, Resources, Writing - Review and Editing, Supervision

LAO - Conceptualization, Resources, Writing - Review and Editing, Supervision, Funding Acquisition

## 10. SUPPLEMENTARY MATERIAL

**Figure S1:**
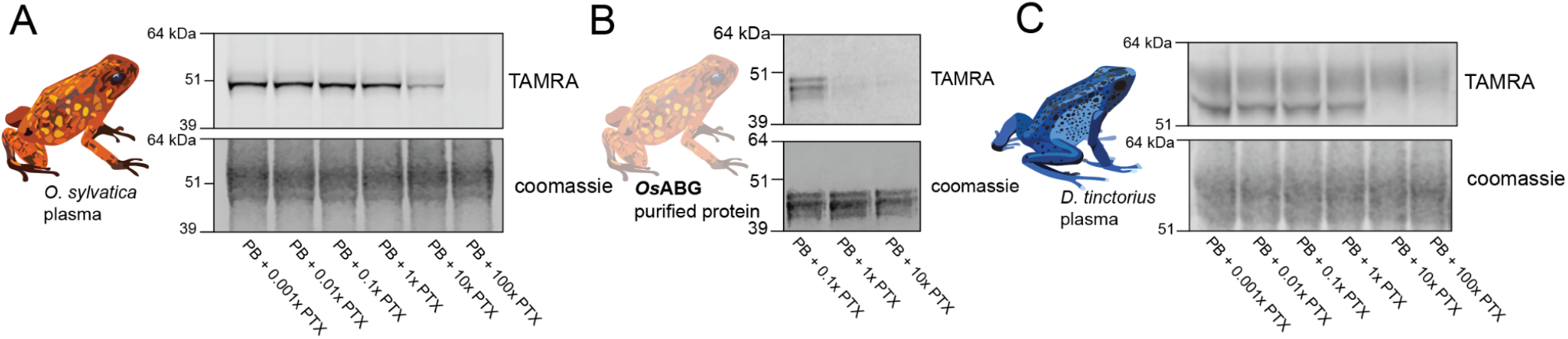
Dose response of photoprobe competition. **(A)** Plasma from *O. sylvatica* shows crosslinking to the photoprobe, and competition by pumiliotoxin **251D** (PTX) occurs when there is 10:1 PTX:photoprobe concentration in the reaction. **(B)** Purified O. sylvatica Alkaloid Binding Globulin (ABG) shows crosslinking to the photoprobe and competition by PTX when there is a 1:1 PTX:photoprobe concentration. **(C)** Plasma from *D. tinctorius* shows crosslinking to the photoprobe, and competition by PTX occurs when there is 10:1 PTX:photoprobe concentration in the reaction.

**Figure S2:**
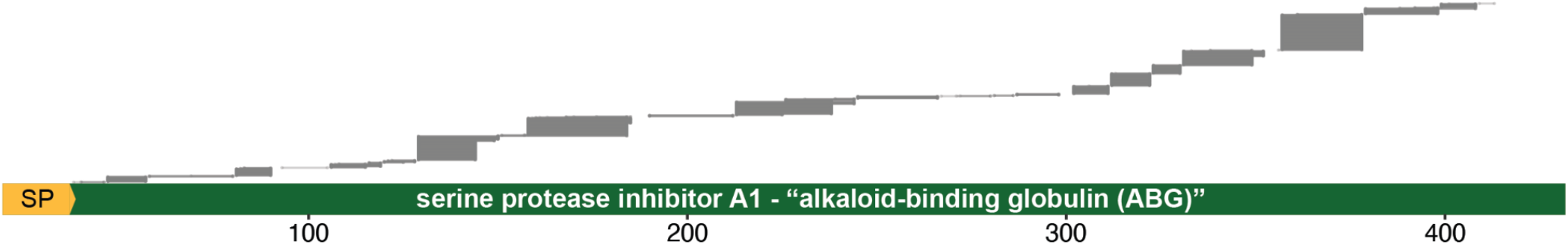
Peptide coverage over ABG protein sequence. All unique peptides from one replicate of the proteomics were mapped onto the protein sequence of *O. sylvatica* ABG, showing that there were peptides covering the full length of the protein.

**Figure S3:**
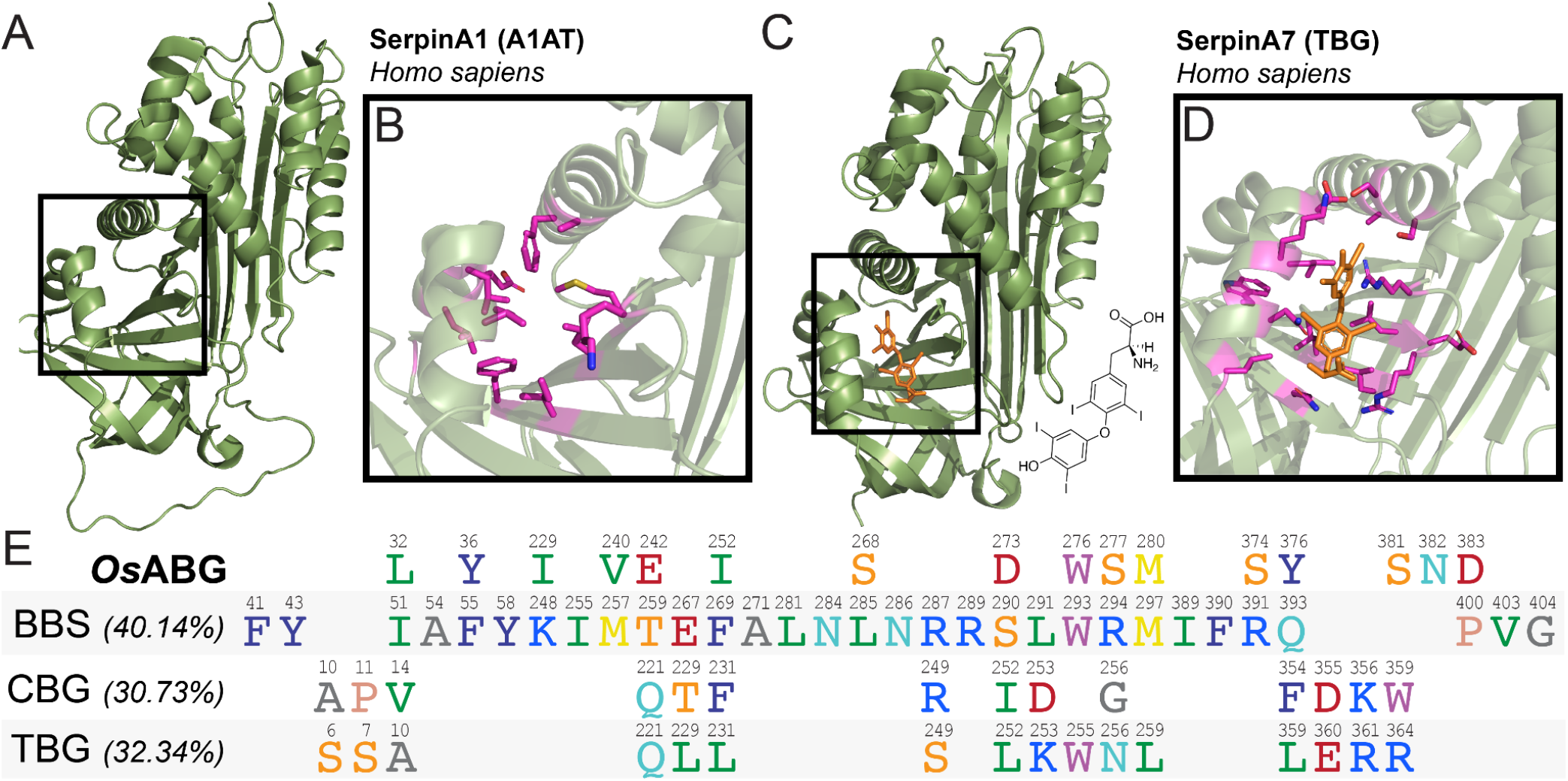
Structure and binding pockets of closely related serpinA proteins. **(A)** Crystal structure for human SerpinA1/alpha-1-antitrypsin (PDB# 1HP7) contains the structural elements of other serpin binding pockets (black box), however is not documented to have small-molecule binding capabilities. **(B)** Close-up of A1AT (PDB# 1HP7), with pocket residues highlighted in magenta. **(C)** Crystal structure for human SerpinA7/thyroxine-binding globulin (PDB# 2RIW) binds thyroxine (orange) in the same structural pocket as other serpins. **(D)** Close-up of thyroxine binding pocket of TBG (PDB# 2RIW), with proximal residues highlighted in magenta. Thyroxine structure is displayed on the top right. **(E)** Alignment of proximal residues (within 5 angstroms of small molecule) across small molecule binding serpins *Os*ABG, biliverdin binding serpin (BBS, PDB# 7RBW), corticosteroid binding globulin (CBG, PDB# 2V95), and thyroxine binding globulin (TBG, PDB# 2RIW) shows that most residues that may be involved in coordinating small molecule binding are not conserved, despite the structural conservation of the binding pocket. Percentages indicate the total percent identity of the protein sequences, the small number above each residue indicates the position of that amino acid in the protein sequence. Only proximal residues are shown, blank spaces are not representative of any specific sequence or spacing.

**Figure S4:**
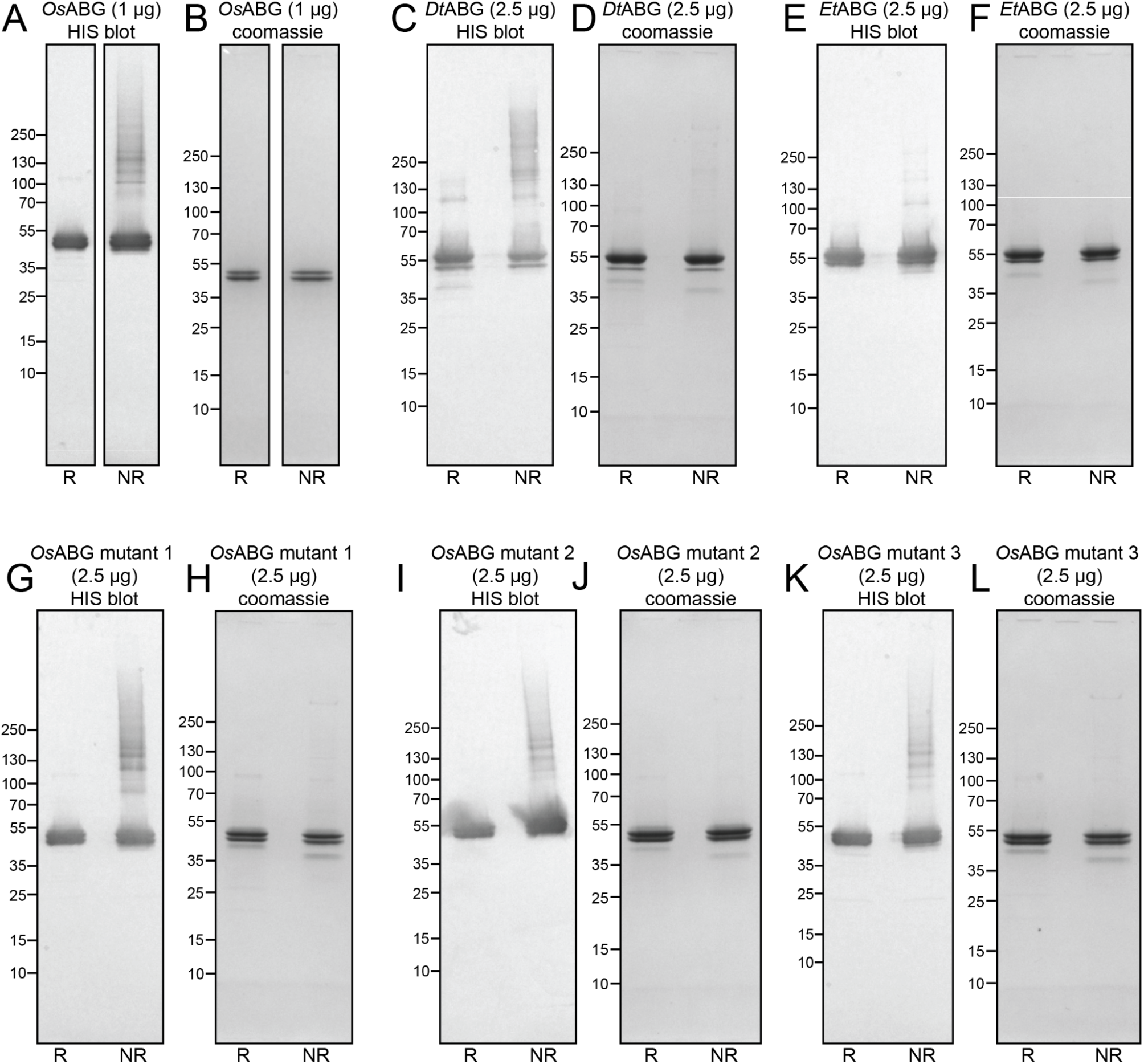
Recombinant expression and purification of Alkaloid Binding Globulin (ABG) proteins in insect cells. HIS blot **(A)** and coomassie **(B)** of the recombinantly expressed and purified *O. sylvatica* alkaloid-binding globulin (*Os*ABG) show a clear doublet pattern in both reduced (R) and non-reduced (NR) conditions. HIS blot **(C)** and coomassie **(D)** of recombinantly expressed *D. tinctorius* ABG, identified by the most-homologous protein to OsABG in the *D. tinctorius* transcriptome, show a clear purification. HIS blot **(E)** and coomassie **(F)** of recombinantly expressed *E. tricolor* ABG, identified by the most-homologous protein to OsABG in the *E. tricolor* transcriptome, show a clear purification. HIS blot **(G)** and coomassie **(H)** of recombinantly expressed *Os*ABG mutant 1 (Y36A, W276A, S374A, D383A) shows a clear purification. HIS blot **(I)** and coomassie **(J)** of recombinantly expressed *Os*ABG mutant 2 (Y36A, S268A, D273A, D383A) shows a clear purification. HIS blot **(K)** and coomassie **(L)** of recombinantly expressed *Os*ABG mutant 3 (D383A) shows a clear purification.

